# Beyond urbanization metrics: Using graphical causal models to investigate mechanisms in urban ecology and evolution

**DOI:** 10.1101/2024.09.03.611045

**Authors:** Jesse B. Borden, James P. Gibbs, John P. Vanek, Bradley J. Cosentino

**Affiliations:** Department of Biology, Hobart and William Smith Colleges, 300 Pulteney Street, Geneva, NY, 14456, USA; Department of Environmental Biology, The State University of New York College of Environmental Science and Forestry, 1 Forestry Drive, Syracuse, NY 13210, USA; New York Natural Heritage Program, The State University of New York College of Environmental Science and Forestry, 625 Broadway, Albany, NY 12207, USA

**Keywords:** causal inference, directed acyclic graphs, eastern gray squirrel, *Sciurus carolinensis*, urban ecology, urban evolution

## Abstract

As the fastest-growing form of land use, urbanization drives profound environmental change that reshapes biodiversity. Studies often characterize biodiversity patterns along urban-rural gradients with statistical models that include one or more generic indices of urbanization. Such models may be useful for prediction, but they do not permit explicit tests of causal hypotheses by which urbanization mediates ecological and evolutionary processes. Here, we show how a graphical causal modeling framework with directed acyclic graphs (DAGs) can be used to design clear conceptual models and inform appropriate statistical analysis to better evaluate mechanistic hypotheses about the effects of urbanization on biodiversity. We first introduce the basic structure of DAGs and illustrate their value with simulated datasets. We then apply the framework to a case study on coat color variation in eastern gray squirrels (*Sciurus carolinensis*) along an urbanization gradient in Syracuse, New York, USA. We show how statistical models ungrounded in causal assumptions are difficult to interpret and can lead to misleading conclusions about mechanisms in urban ecology and evolution. In contrast, DAGs – by making causal assumptions transparent – help researchers identify appropriate control variables for statistical models to estimate the effects of interest. When applied to our case study, statistical models informed by a DAG revealed a surprising finding: although squirrel melanism was more prevalent in urban than rural populations, the prevalence of melanism was constrained by components of environmental change common to cities, namely roads, forest loss, and predator activity, contrary to expectations. Managing biodiversity in an increasingly urbanized world will require a mechanistic understanding of how urbanization impacts biodiversity patterns; graphical causal models such as DAGs can provide a powerful approach to do so.

## Introduction

Urban areas represent the fastest growing land use type and have profound effects on biodiversity (Aronson et al. 2014; Ritchie and Roser 2018; Szabó et al. 2023). Urbanization consists of a suite of abiotic and biotic changes driven by high concentrations of humans and the physical environment they build (Ritchie and Roser 2018). The ecological and evolutionary consequences of urbanization are often studied using the urban-rural gradient approach, in which components of biodiversity are compared along a landscape-scale sampling area from the urban core to rural habitats (McDonnell and Pickett 1990; McDonnell and Hahs 2008). The urban-rural gradient approach has been used to document associations of population size, species distributions, community composition, and genetic and phenotypic trait variation with urbanization (Winchell et al. 2016; Moll et al. 2019; Cosentino and Gibbs 2022; Santangelo et al. 2022).

Although the urban-rural gradient method has been helpful to describe spatial patterns of biodiversity, reliance on univariate metrics of urbanization (e.g., impervious cover) can be problematic when the research goal is to identify processes underlying those patterns (Moll et al. 2019, Poisson et al. 2024). Moreover, the use of statistical models with multiple covariates (e.g., multiple linear regression) does not guarantee valid causal inference. Interpreting parameter estimates from statistical models as estimated causal effects requires explicit assumptions about the causal relationships among covariates in the model. In many studies those relationships are unstated, and covariates are included in models without regard to the causal connections between them (McElreath 2020).

Understanding mechanisms by which urbanization generates patterns in ecology and evolution is complicated by at least two factors. First, environmental change associated with urbanization can affect biodiversity through multiple causal pathways (Alberti et al. 2020). For example, road networks can *directly* affect biodiversity through vehicular collisions and noise pollution (Francis et al. 2012; Bennett 2017; Kok et al. 2023), but roads also have *indirect* effects by contributing to habitat fragmentation and pollution runoff, altering predator-prey dynamics, or facilitating the spread of invasive species (Bowman et al. 2010; Bennett 2017; Mumma et al. 2019; Cerqueira et al. 2021). When a univariate model is used to test how a biodiversity component is related to road cover, the estimated effect combines the direct and indirect effects. In some cases, these effects can offset one another, obscuring different causal mechanisms and leading to the conclusion that road cover has no impact.

Second, many environmental conditions covary along urbanization gradients – for example, road extent with housing density or light pollution – confounding the interpretation of the effect of any one axis of environmental variation (Moll et al. 2019). Including multiple covariates in a regression model without a causal rationale can introduce bias. For example, if light level mediates the effect of road cover on biodiversity, including light as a covariate blocks part of the causal pathway and underestimates the total effect of road cover. Conversely, failing to include confounding variables that affect both road cover and biodiversity can lead to spurious associations. Statistical models designed without regard to causal assumptions often yield parameter estimates that conflate true causal effects with noncausal confounding, rendering a mechanistic interpretation nearly impossible.

One way to improve causal inference is to use *graphical causal models*, which have been shown effective across the sciences (e.g., Rohrer 2018; McElreath 2020; Arif and MacNeil 2023; Arif and Massey 2023). Our goal is to show how these models can be leveraged to strengthen inferences about urbanization effects on biodiversity. Specifically, we highlight one type of graphical causal model, the *directed acyclic graph* (DAG), as an approach for specifying causal assumptions about urban systems, including direct effects, indirect effects via mediators, and confounding paths. We first provide a basic introduction to DAGs in the context of urban ecology and evolution, using simulated datasets to show how the causal meaning of statistical estimates depends on assumptions specified in the DAG. We then apply the method to a case study in urban evolution, building a DAG to represent our causal assumptions and guide statistical models to test hypotheses about urbanization effects on coat color variation in eastern gray squirrels (*Sciurus carolinensis*).

## Directed acyclic graphs for urbanization effects

Directed acyclic graphs are visual diagrams representing hypothesized causal relationships among variables. DAGs help clarify causal assumptions and inform decisions about study design and statistical analysis when the research goal is to understand mechanisms driving observed patterns (Arif and Massey 2023). DAGs consist of variables connected by arrows, with each arrow representing a direct causal effect. Importantly, causal pathways in DAGs are probabilistic: the path *X* → *Y* means *X* influences the probability distribution of *Y*, not that *Y* is a deterministic function of *X*.

Consider a hypothetical example, in which an urban ecologist is interested in the effects of housing density on bird species diversity. We created an example DAG that might represent a researcher’s causal assumptions related to the research question (Fig. 1). The DAG shows housing density is expected to affect bird diversity via two pathways: 1) the **direct effect** *Housing density* → *Species diversity*, perhaps representing the expectation that housing structures can provide nesting habitat for some bird species, and 2) the **indirect effect** *Housing density* → *Green space* → *Species diversity*, which might reflect an ecologist’s expectation that housing structures reduce green space, while more green space increases species diversity. Here, green space functions as a **mediator**, transmitting an effect of housing density on species diversity. The **total effect** of a variable on an outcome is simply the sum of its direct and indirect effects. Thus, based on our example DAG, the total effect of housing density includes its direct effect plus the effect mediated by green space.

**Fig. 1.**
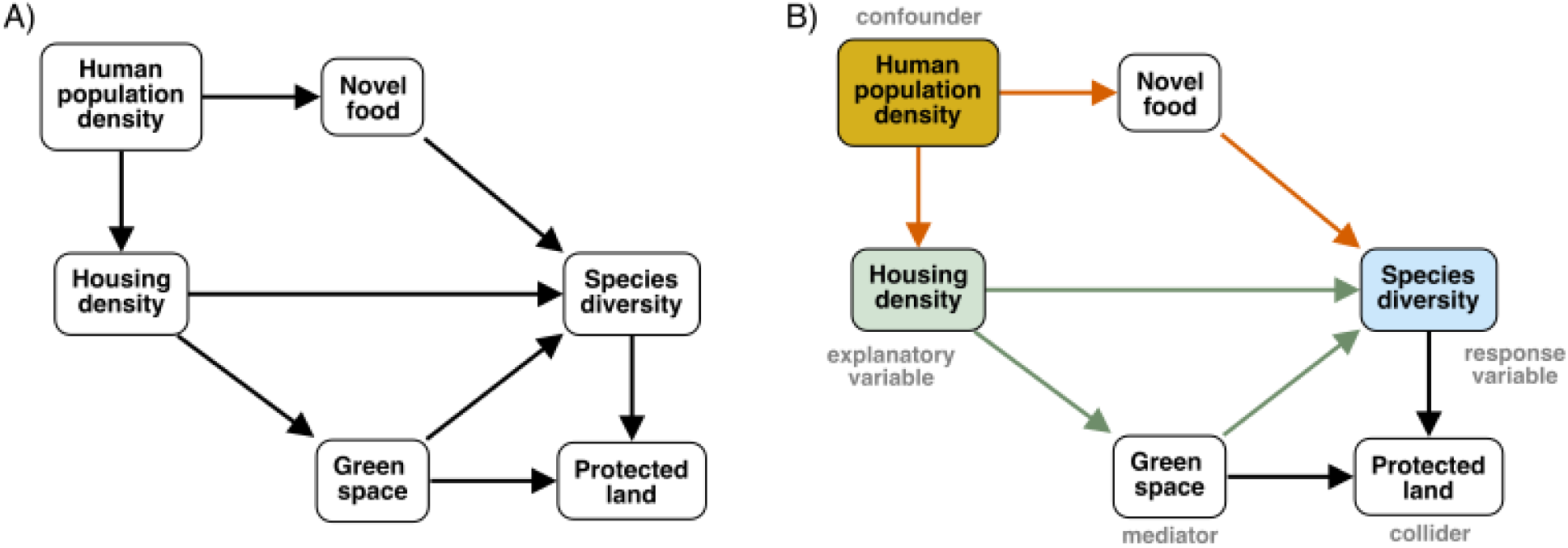
Hypothetical directed acyclic graph (DAG) for a study of the causal effects of housing density on bird species diversity. Boxes are measured variables, and arrows represent assumed direct causal effects between variable pairs. Panels include an unlabeled version (A) and a labeled version (B) identifying the explanatory (green box) and response (blue box) variables of interest, the causal pathways of interest (green arrows), and important causal structures to identify when estimating the causal pathways of interest. Estimating effects of housing density on species diversity requires adjusting for the confounding effect of human population density (orange box) that opens a backdoor path (orange arrows). In contrast, protected land is a collider and does not cause bias unless conditioned on. Estimation of the total effect of housing density requires adjusting for the confounder only, whereas estimation of the direct effect requires adjusting for the confounder and the mediator (green space).

Two additional causal structures are essential to identify in DAGs: confounders and colliders. In our example DAG, the variables human population density and novel food are on a **backdoor path** connecting housing density and species diversity: *Housing density* ← *Human population density* → *Novel food* → *Species diversity* (Fig. 1). This path might reflect an ecologist’s expectation that human population density increases both housing density and novel food (e.g., bird feeders), and that novel food increases species diversity. A backdoor path is any non-causal pathway involving a variable *Z* that causally affects an explanatory variable *X* and the response variable *Y* (directly or indirectly), having the general structure *X* ← *Z* → *Y*. In this case, *Z* is a **confounder** because it affects both *X* and *Y*, which can create a spurious association or mask a true causal relationship between them. When considering the effect of housing density on species diversity, human population density is a confounder because it affects housing density directly and species diversity indirectly via novel food.

Another important causal structure in the example DAG involves the variable protected land: *Green space* → *Protected land* ← *Species diversity* (Fig. 1). Here, the amount of protected land is influenced by green space and species diversity, perhaps reflecting an ecologist’s expectation that protected status is most likely to be prioritized for locations with high species diversity or extensive green space. In this example, protected land is a **collider**. A collider has the form *X* →*Z* ←*Y*, where the collider *Z* is affected by *X* and *Y*.

The causal structures highlighted here are essential to identify when using statistical models to estimate causal effects. To correctly estimate the effects of interest, researchers need to parameterize statistical models in a specific way based on causal assumptions expressed in the DAG. For example, to correctly estimate the direct and total effects of housing density on species diversity, it is necessary to adjust for the confounding pathway, typically by including human population density or novel food (or both) as covariates in a regression model. But care must be taken to decide which covariates to include in a model. If the direct effect of housing density was of interest, one needs to adjust not only for the confounding effect of human population density, but also the mediating effect of green space. In contrast, green space should *not* be included as a covariate when estimating the total effect of housing density, as it functions as a mediator of housing density and therefore contributes to the total effect. Similarly, colliders do not cause bias by default and should not be included as a covariate when they are on a path from the explanatory variable to the response variable. Including protected land in a regression model with housing density would bias estimates of the housing density effects.

We used a data simulation exercise to explicitly demonstrate how valid estimation of causal effects with statistical models depends on causal assumptions in a DAG. We first used our example DAG (Fig. 1) to generate synthetic datasets in R (R Core Team 2024). On their own, DAGs do not imply a functional form of causal relationships between variables (e.g., positive-negative, linear-nonlinear, additive-interactive). For the simulation, we assumed all relationships were linear and additive, and we defined the direction and strength of each direct effect in the DAG as standardized regression coefficients (Table 1). We then used the assumed effects to simulate data for 100 sampling locations, with each variable standardized to mean = 0 and SD = 1. All effects were probabilistic, such that the observed value of each variable at each sampling location was a function of its causal ancestors plus a random error term drawn from a normal distribution with mean = 0 and SD = 0.5. We generated a total of 1000 simulated datasets.

**Table 1.** Standardized regression coefficients representing the direction and magnitude of direct effects in our hypothetical DAG (Fig. 1) used to simulate observed datasets.

| *Explanatory variable* | *Response variable* | *Standardized coefficient* |
| --- | --- | --- |
| Human population density | Novel food | 0.9 |
| Human population density | Housing density | 0.8 |
| Housing density | Green space | -1.0 |
| Housing density | Species diversity | 0.3 |
| Novel food | Species diversity | 0.4 |
| Green space | Species diversity | 0.8 |
| Green space | Protected land | 1.0 |
| Species diversity | Protected land | 0.8 |

For each simulated dataset, we fit statistical models to estimate the effects of housing density on species diversity. We used the *dagitty* package (Textor et al. 2016) to identify appropriate **adjustment sets** to estimate the direct and total effects of housing density on species diversity. An adjustment set consists of the covariates from a DAG that should be included in a statistical model to produce unbiased estimates of a causal effect of interest (Textor and Liśkiewicz 2011). There are often multiple possible adjustment sets for a given causal effect, and in our example, the algorithm used by *dagitty* identified two valid adjustment sets for the direct effect and two valid adjustment sets for the total effect. For each simulated dataset, we fit a regression model corresponding to each adjustment set. For comparison, we fit two additional models commonly used in the literature: 1) a univariate model with housing density as the only predictor of species diversity, and 2) a full model including all covariates.

The simulation results showed the total effect of housing density on species diversity was accurately estimated when adjusting for the confounding effect of human population density, which required including either human population density or novel food in the regression model (Fig. 2A, B). The direct effect of housing density was accurately estimated when adjusting for the same confounding pathway, as well as the mediating effect of green space (Fig. 2C, D). In contrast, the univariate model and full model both produced biased estimates of the housing density effects (Fig. 2E, F). The univariate model conflates the direct and indirect effects of housing density with the noncausal associations with human population density and novel food, leading to a biased estimate that recovers neither of the true effects. The full model approximated the direct effect of housing density because it adjusts for the confounder (human population density) and mediator (green space), but it underestimated the direct effect because it also included the collider, protected land. Overall, the data simulation illustrates how the meaning of an estimated regression coefficient depends on both the assumed causal relationships among variables specified in a DAG *and* the specific parameterization of a statistical model.

**Fig 2.**
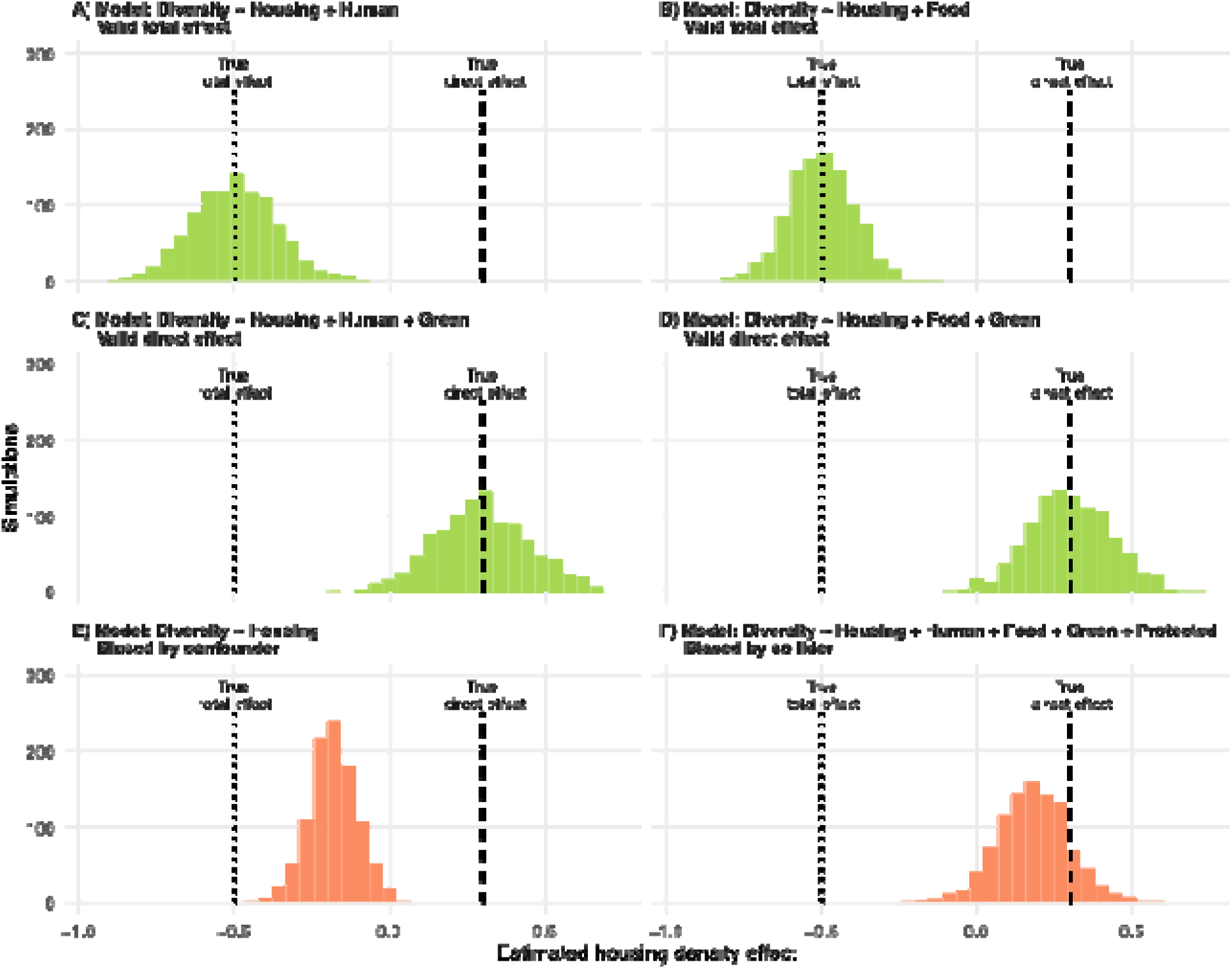
Histograms of the estimated housing density coefficient from regression models fit to 1000 simulated datasets generated from our example DAG (Fig. 1, Table 1). Each panel (A-F) shows the distribution of the estimated housing density coefficient from a single regression model. Model parameterization is shown as a formula with species diversity (Diversity) as a function of (∼) housing density (Housing) and other covariates: human population density (Human), novel food (Food), green space (Green), and protected land (Protected). Vertical dashed lines show the true total and direct effects of housing density on species diversity. Histogram colors distinguish models that produce accurate estimates of the causal effect (green) from models that produce biased effects (orange).

DAGs are useful to make causal assumptions clear and to inform statistical analysis, but their use does not guarantee valid causal inference. A critical assumption when using a DAG is that it specifies the correct causal structure of a system. In practice, researchers should apply prior knowledge of their study system and consider whether non-trivial causes are missing from a DAG, especially variables that lie on confounding pathways between the explanatory and response variables of interest. Importantly, confounding effects should be included in DAGs even when they involve unmeasurable variables, as there are approaches to address unobserved confounders (Byrnes and Dee 2025).

At the same time, not all variables need to be included in a DAG. Nearly all causal pathways can be decomposed into more detailed steps (Shipley 2016), and, like any model, DAGs are simplifications of reality. If a variable functions solely as a mediator between two variables, it can often be removed from the DAG without changing the implied causal structure. Objections may be raised about omitted variables, but we argue that debate about the structure of DAGs is exactly what makes them so valuable. By forcing researchers to state their causal assumptions explicitly, DAGs promote transparency, constructive criticism, and ideally stronger inferences.

## Case study: applying DAGs to test causal hypotheses about coat color variation in eastern gray squirrels

### Study species and hypothesized urbanization effects

Eastern gray squirrels (*Sciurus carolinensis*) provide an excellent study species to show how DAGs can be used to help disentangle mechanisms by which urbanization affects ecological and evolutionary outcomes. Eastern gray squirrels are common across urban and rural landscapes in their native range of eastern North America. Individuals tend to have one of two coat color morphs: gray or black (melanic) inherited in a simple Mendelian fashion (McRobie et al. 2009).

The prevalence of melanism tends to be greater in cities than adjacent rural forests (Gibbs et al. 2019; Cosentino and Gibbs 2022; Cosentino et al. 2023). Multiple hypotheses have been proposed about the ecological and evolutionary processes maintaining these urban-rural clines (Cosentino and Gibbs 2022; Cosentino et al. 2023). For example, the melanic morph is visually more conspicuous than the gray morph, such that predation may select for the more cryptic gray morph (Proctor et al. 2025). Thus, both predator activity and habitat context (e.g., sources of refuge from predators, including trees and buildings) could alter the prevalence of melanism.

Additionally, road mortality has been hypothesized to maintain clines in melanism, as the melanic morph is more visible to humans and underrepresented among roadkill compared to the gray morph (Gibbs et al. 2019; Proctor et al. 2025; Parlin et al. 2025).

### Generating an initial DAG

We created a DAG to represent our causal assumptions for how predator activity, components of habitat structure that affect refuge availability (building cover, forest cover, and forest fragmentation), and road cover affect spatial variation in the prevalence of melanism along an urbanization gradient (Fig. 3A). We expected melanism would be greatest in areas of refugia from predators (high building cover, high forest cover, low forest fragmentation) and high road cover, and lowest in areas of high predator activity. Critically, while these variables are all hypothesized to affect the prevalence of melanism directly (Table 2), they can covary along urban-rural gradients and confound one another, highlighting the necessity of using a DAG to identify potential confounds and appropriate adjustments to estimate the effects of interest. For example, forest cover and forest fragmentation may directly affect the prevalence of melanism by altering crypsis or availability of refuge, but such habitat alteration could also indirectly affect the prevalence of melanism by changing predator activity (Bateman et al. 2012).

**Fig. 3.**
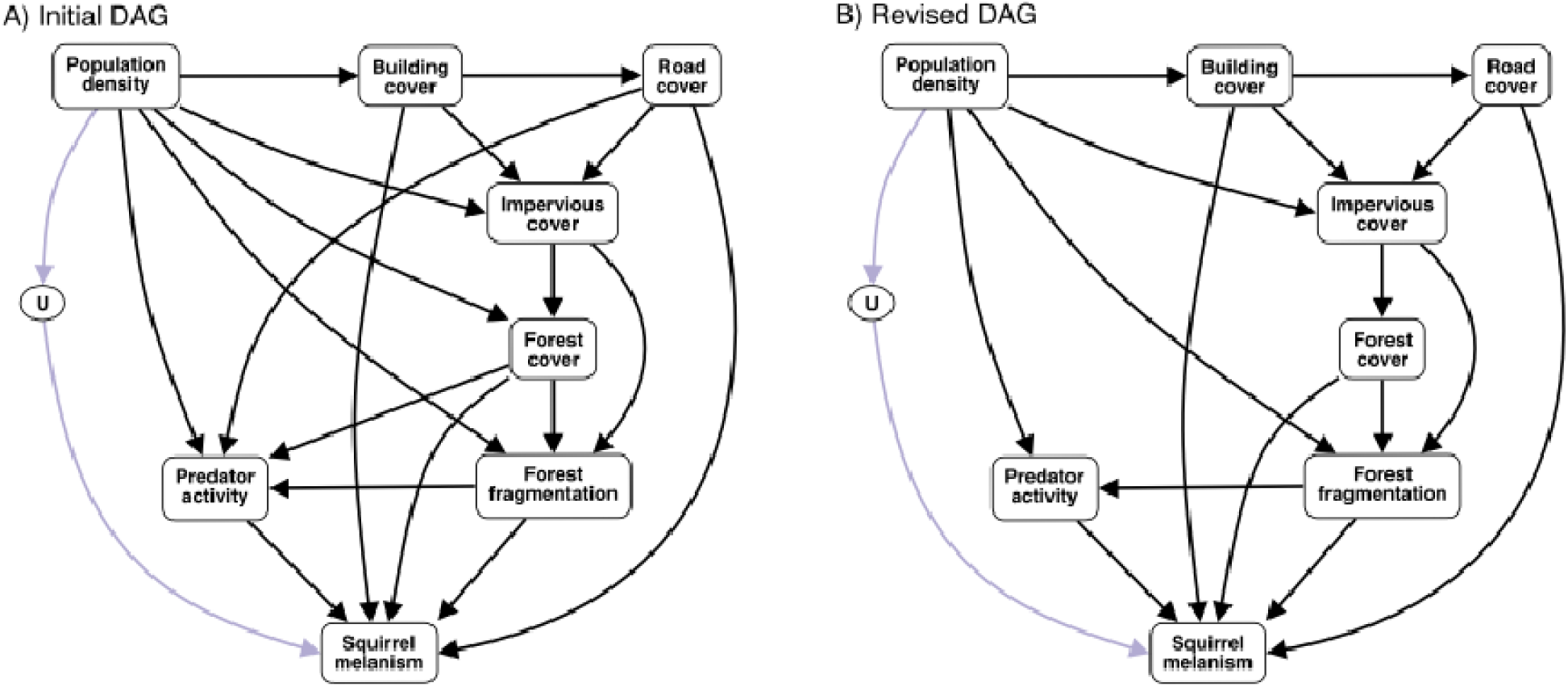
Directed acyclic graphs (DAG) representing our scientific model, showing hypothesized causal relationships between variables that could affect variation in melanism in eastern gray squirrels (*Sciurus carolinensis*) along urbanization gradients. An initial DAG was generated representing our original hypotheses (A), and the DAG was revised by dropping direct effects among explanatory variables that were not empirically supported (B) (see *Statistical analysis* below). Arrows indicate hypothesized direct effects. Boxes represent measured variables, and circles represent unobserved variables. “U” represents unobserved effects of people on the prevalence of squirrel melanism, including translocations and hunting pressure, which are known to occur but remain unmeasured. Arrow color distinguishes pathways that were estimable with measured variables (black) from pathways that were not estimable due to unobserved processes (purple).

**Table 2.** Environmental variables hypothesized to affect the prevalence of melanism in eastern gray squirrels (*Sciurus carolinensis*) and data used to measure those variables.

| Environmental variable | Rationale | Hypothesized Direction | References | Variable data source |
| --- | --- | --- | --- | --- |
| Human population density | The foundational driver of urbanization, increasing building and road cover, driving deforestation and habitat fragmentation, and causing many unmeasured changes. | (+) | (Glover and Simon 1975) | NASA's CIESIN gridded human population dataset (30 arcsec; Center For International Earth Science Information Network-CIESIN-Columbia University 2017) |
| Building cover | Alters habitat structure and availability by reducing tree cover, but provides nesting sites, novel microclimates, and shading that may be important for crypsis. | (+) | (Benson 2013; Parker and Nilon 2008) | Cornell University Geospatial Information Repository |
| Road cover | Roadkill is a major driver of squirrel mortality in urban contexts particularly roads with $\geq 30$ mph speed limit. Melanic individuals are more visible to drivers on roads, and they have lower mortality rates from vehicular collisions than the gray morph. | (+) | (McCleery et al. 2008; Gibbs et al. 2019; Parlin et al. 2025; Proctor et al. 2025) | Cover of roads with at least 30 mph speed limit from NYS GIS Resources repository for public data (https://data.gis.ny.gov) |
| Forest cover | Increases resource availability, including food, nests, and refuge from predators. | (+) | (Parker and Nilon 2008; Koprowski et al. 2016; Merrick et al. 2016) | ESA WorldCover dataset derived from Sentinel-1 and Sentinel-2 satellite imagery (10 m resolution; Zanaga et al. 2022) |
| Forest fragmentation | Restricts movements and may affect fitness of color morphs differently by altering predator activity and crypsis of each morph assuming fragmented areas are more visually open. | (-) | (Parker and Nilon 2008; Koprowski et al. 2016) | Measured as disjunct core area density based on forest cover from ESA WorldCover dataset derived from Sentinel-1 and Sentinel-2 satellite imagery (Zanaga et al. 2022) |
| Mammalian predator activity | Mammalian predator activity alters predation risk as well as the landscape of fear. Differences in crypsis or behavior may cause morphs to be affected differently by predator activity. | (-) | (Gibbs et al. 2019; Cosentino and Gibbs 2022; Proctor et al. 2025) | Quantified from camera trap detections |

The foundation of our DAG was human population density, operating on the assumption that environmental change along urbanization gradients is ultimately driven by people. We included a direct effect of human population density on building cover (Glover and Simon 1975). Because roads are the main way to access buildings, we included a direct effect of building cover on road cover. Buildings and roads were assumed to be direct causes of impervious cover, and we included a direct effect of human population density on impervious cover to represent other types of impervious surface that people build (e.g., sidewalks, parking lots). We included a direct effect of impervious cover on forest cover, as the creation of infrastructure requires tree removal (Liu et al. 2014). Forest cover extent is known to decrease forest fragmentation (Fahrig 2003, 2017), so we included a direct effect of forest cover on fragmentation. We also included a direct effect of impervious cover on forest fragmentation, as landscape elements like buildings and roads cause discontinuities in forest cover, regardless of their effects on the absolute amount of forest (Liu et al. 2014; Liu et al. 2019). Human population density was included as a direct effect of forest cover and fragmentation to account for other ways that people remove and fragment forested areas (e.g., gardening and open parks). We included direct effects of road cover, forest cover, and fragmentation on mammalian predator activity, as all three have been linked to urban carnivore distribution and behavior (Ordeñana et al. 2010; Bateman et al. 2012).

Although DAGs are useful for identifying and adjusting for confounders, unobserved or unmeasured confounders still create the potential for bias (Byrnes and Dee 2025). For example, the high prevalence of melanism among squirrels in some cities may be due in part to historical introductions by people (i.e., propagule pressure) or reduced hunting pressure in cities (Cosentino et al. 2023), variables that were not possible to measure. Including unobserved variables like these in a DAG can be important to make gaps in knowledge explicit and identify confounding with other measured variables. As such, we added an unobserved variable to our DAG to represent unmeasured mechanisms by which people could mediate coat color change (“U”). We included a direct effect of human population density on U and a direct effect of U on the prevalence of melanism (Fig. 3A). By including this unobserved variable in the DAG (Fig. 3A), we were able to identify how backdoor paths which include U as a confounder could be closed by adjusting for human population density (Table 3).

**Table 3.** Posterior means, 95% credible intervals, and adjustment sets for parameter estimates of direct effects, total effects, and univariate models for predictors of coat color (proportion melanic) among eastern gray squirrels (*Sciurus carolinensis*). Separate models were fit to estimate direct and total effects of predictor variables on proportion melanic with appropriate adjustment covariates based on our directed acyclic graph (Fig. 3B). Univariate estimates were based on models with only the predictor variable and no adjustment covariates. Predictor variables included human population density (Human), building cover (Building), road cover (Road), impervious cover (Impervious), forest cover (Forest), forest fragmentation (Frag), and mammalian predator activity (Predator).

| **Direct Effects** |  |  |  |  |
| --- | --- | --- | --- | --- |
| *Predictor* | *Mean* | *LCL* | *UCL* | *Adjustments* |
| Human | - | - | - | - |
| Building | 0.02 | -0.29 | 0.32 | Human, Road, Forest, Frag |
| **Road** | **-0.69** | **-1.01** | **-0.39** | Human, Building, Forest, Frag |
| Impervious | - | - | - | - |
| **Forest** | **0.47** | **0.17** | **0.78** | Human, Building, Road, Frag |
| Frag | 0.28 | -0.07 | 0.64 | Human, Impervious, Forest, Predator |
| **Predator** | **-0.54** | **-0.83** | **-0.26** | Human, Frag |
| **Total Effects** |  |  |  |  |
| *Predictor* | *Mean* | *LCL* | *UCL* | *Adjustments* |
| **Human** | **0.42** | **0.18** | **0.67** | - |
| **Building** | **-0.33** | **-0.54** | **-0.10** | Human |
| **Road** | **-0.28** | **-0.54** | **-0.01** | Building |
| **Impervious** | **-0.80** | **-1.42** | **-0.19** | Human, Building, Road |
| Forest | 0.16 | -0.08 | 0.42 | Impervious |
| **Frag** | **0.41** | **0.07** | **0.74** | Human, Impervious, Forest |
| **Predator** | **-0.54** | **-0.83** | **-0.26** | Human, Frag |
| **Univariate Estimates** |  |  |  |  |
| *Predictor* | *Mean* | *LCL* | *UCL* | *Adjustments* |
| **Human** | **0.42** | **0.18** | **0.67** | - |
| Building | -0.04 | -0.22 | 0.15 | - |
| Road | -0.15 | -0.34 | 0.02 | - |
| Impervious | -0.15 | -0.35 | 0.04 | - |
| Forest | 0.20 | -0.01 | 0.42 | - |
| Frag | 0.15 | -0.05 | 0.35 | - |
| Predator | -0.14 | -0.39 | 0.12 | - |

### Study area and sampling

To parameterize the DAG, we used data on coat color variation in eastern gray squirrels across an urban-rural gradient in Syracuse, New York, USA (43.0469° N, -76.1444° W). Syracuse had an estimated human population of 146,000 (in 2022) and occurs in a humid continental climate (Köppen 1936) with four distinct seasons, including hot summers and cold winters. Tree cover ranges from 4.5% to 47% across the city, averaging 27% (Nowak et al. 2016). The surrounding rural landscape is a mosaic of suburban areas, small villages, agricultural lands, wetlands, and woodlands (Supplemental Fig. S1).

We used camera traps and point count surveys to estimate prevalence of color morphs at 49 sites across the urban-rural gradient between September 2021 and October 2022 (Supplemental Fig. S1; Cosentino et al. 2023). Sites were selected in urban greenspaces and rural woodlots with mature deciduous trees, spanning the urban-rural gradient (range = 1.3–23.7 km from the city center) and separated by at least 1 km. A single camera trap (Browning Strikeforce Pro XD) was fastened to a tree 30-50 cm off the ground at each site and activated for 46-379 days (median = 263 days), and daily detection histories were generated for each color morph. We also conducted 2-7 (median = 5) visual point count surveys at each site, recording the number of squirrels seen of each color morph during a 3-min period. See Cosentino et al. (2023) for detailed field methods.

### Quantifying measured variables

All measured variables in the DAG other than predator activity were measured via remotely sensed datasets with the *raster* and *exactextractr* packages in R (Table 2; Baston et al. 2022; Hijmans et al. 2023). We used a 300-m buffer to summarize variables around each site to encompass a typical home range size of gray squirrels (<5 ha) (Koprowski et al. 2016). Mammalian predator activity was indexed using our camera trap imagery and calculated as the percentage of camera trap days mammalian predators were detected at each site (Bu et al. 2016). Predators detected by camera traps included coyote (*Canis latrans*), gray fox (*Urocyon cinereoargenteus)*, red fox (*Vulpes vulpes*), fisher (*Pekania pennanti*), domestic dog (*Canis familiaris*) and domestic cat (*Felis catus*). Given our primary goal was to illustrate the broader use of DAGs for causal inference, we limited our analysis of predator effects to a general index of activity across mammalian predator species, acknowledging that the effects of mammalian predators on the prevalence of squirrel melanism may be species-specific, and that we are not accounting for avian predation.

### Statistical analyses

We tested hypotheses from our DAG (Fig. 3A) in two stages. First, we used Bayesian generalized linear models to estimate direct effects among measured variables other than squirrel melanism, selecting appropriate distributions for response variables and adjustment covariates based on the DAG using the *dagitty* R package. For example, to estimate the direct effect of building cover on impervious cover, we assumed impervious cover follows a beta distribution and modeled it as a function of building cover, along with road cover as an adjustment. Models were fit with STAN via the R package *brms* (Bürkner 2017) with noninformative priors and standardized predictors. We generated a revised DAG (Fig. 3B) that dropped direct effects when their posterior distributions broadly overlapped 0 (Supplemental Table S1). Next, we estimated the direct and total effects of each variable on squirrel melanism using an integrated hierarchical model combining point count and camera trap data (Kéry and Royle 2020; Cosentino et al. 2023). This model included a process submodel to estimate morph-specific abundances and proportion melanic at each site, and a detection submodel accounting for temperature effects on activity. We estimated direct and total effects of measured variables in our DAG on proportion melanic with linear models, adjusting for DAG-informed covariates as needed. We also fit univariate models with each measured variable as a predictor of melanism to compare inferences with direct and total effects. Hierarchical models were fit in JAGS via the *jagsUI* R package (Plummer 2017, Kellner and Meredith 2024) using mostly noninformative priors, but also with informative priors for detection and abundance intercepts based on prior knowledge. All models were run with four chains and 4000 retained iterations after enough warm-up or burn-in to reach convergence, which was confirmed visually with traceplots and R-hat < 1.01 (Gelman and Hill 2007). Posterior distributions were summarized with means and 95% credible intervals. Details on statistical models are provided in Supplemental Text S1.

## Results and discussion

Human population density had a positive total effect on the prevalence of squirrel melanism, with prevalence of melanism greatest in more urbanized areas (Fig. 4, Table 3). While this aligns with previous studies showing a strong cline along the urbanization gradient (Cosentino and Gibbs 2022; Cosentino et al. 2023), our DAG-informed analyses provided multiple new insights into the potential mechanisms maintaining the urban-rural cline. Remarkably, the total effects of building cover, road cover, and impervious cover (all metrics of human impact commonly used to represent “urbanization effects”) on the prevalence of melanism were each *negative* despite squirrel melanism being most prevalent in the city (Fig. 4, Table 3). These negative total effects suggest the distribution of the melanic morph is limited by various aspects of physical infrastructure, even in more urban areas where melanic individuals are generally most prevalent. Considering the positive total effect of human population density combined with negative total effects of physical infrastructure on squirrel melanism, our results suggest there remain unmeasured cause(s) represented by U in our DAG (Fig. 5) that play an important role in maintaining the urban-rural cline by favoring melanics in the city.

**Fig. 4.**
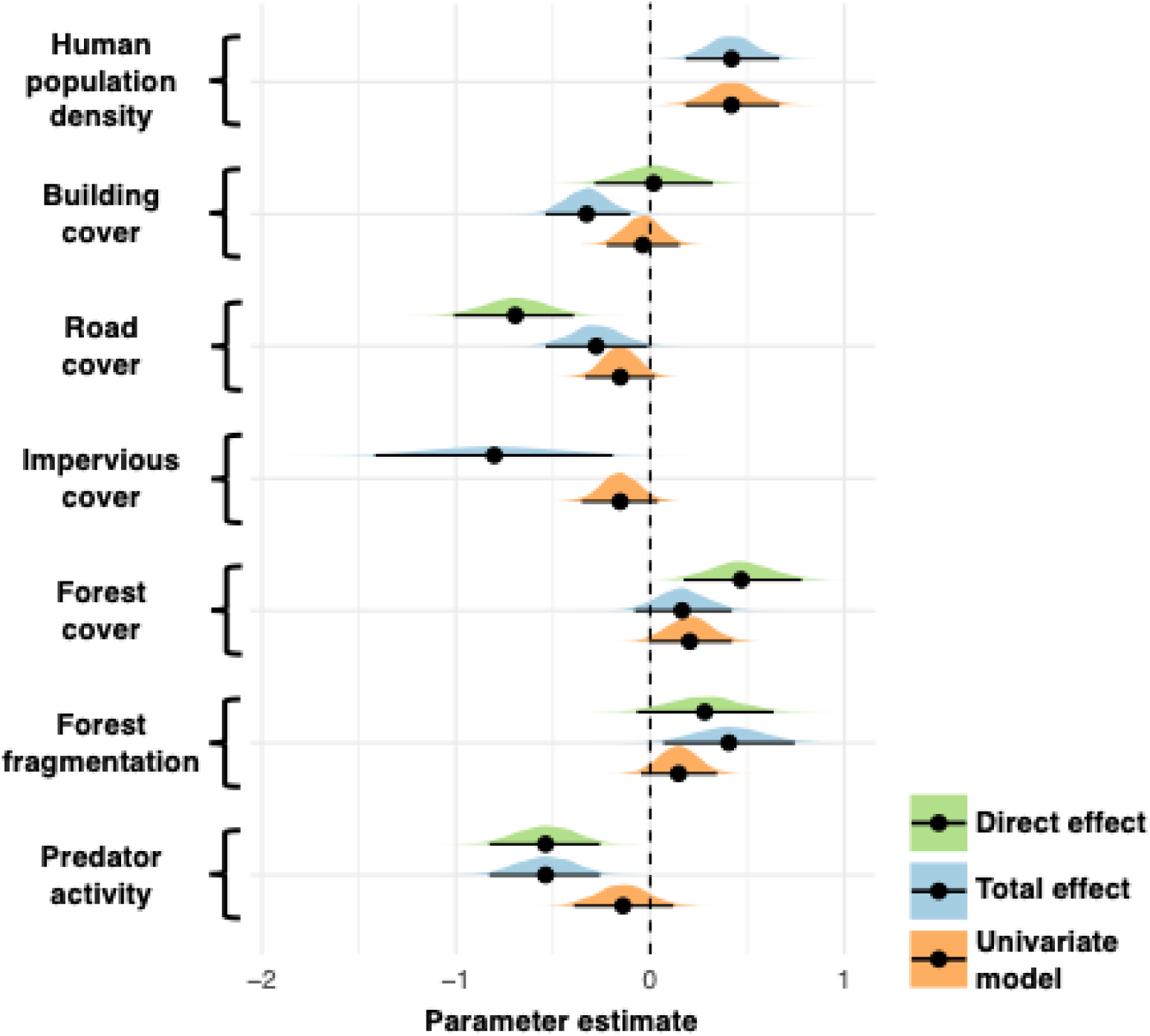
Parameter estimates for direct effects (green), total effects (blue), and univariate models (orange) for predictors of coat color (proportion melanic) among eastern gray squirrels (*Sciurus carolinensis*). The complete posterior distribution is shown for each parameter, along with the mean (black circles) and 95% credible intervals. All parameter estimates were made with standardized predictors. Not all predictors have both direct and total effects based on our directed acyclic graph (Fig. 3B).

**Fig. 5.**
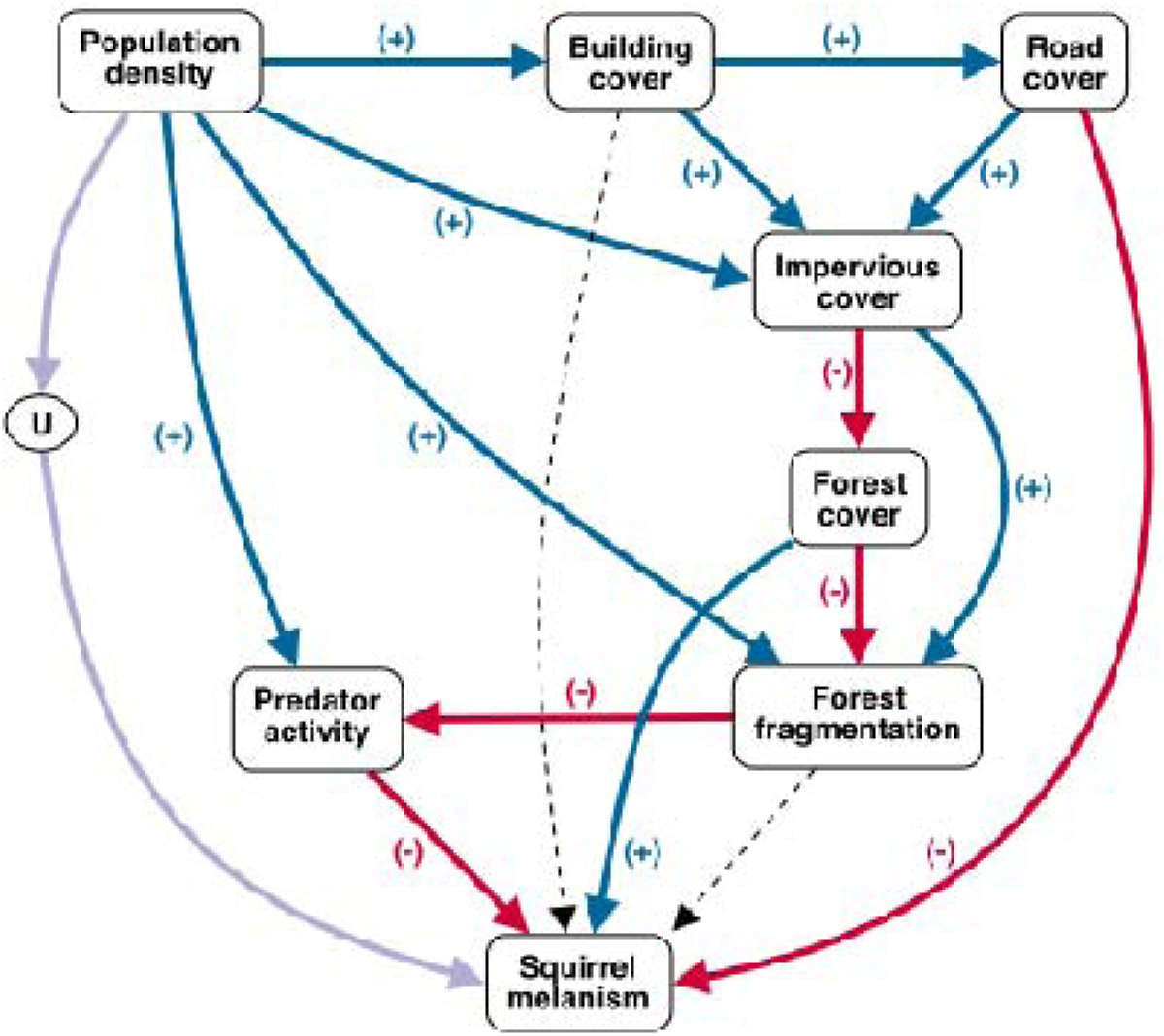
Direction of estimated direct effects overlaid on our directed acyclic graph. Blue solid lines indicate positive direct effects, red solid lines indicate negative direct effects, and black dotted lines indicate effects for which the posterior distribution of the parameter estimate broadly overlapped zero. Purple arrows show causal pathways which could not be estimated due to unobserved processes. Parameter estimates for effects on squirrel melanism can be found in Table 3, and parameter estimates among measured environmental variables can be found in Supplemental Table S1.

The statistical models informed by our DAG provided novel insights into specific mechanisms maintaining the urban-rural cline in melanism. For example, prior research showed melanic individuals are underrepresented among roadkill, relative to the gray morph—a finding attributed to humans more easily detecting and avoiding melanics against gray asphalt when driving (Gibbs et al. 2019, Parlin et al. 2025, Proctor et al. 2025). However, our findings show road cover had strong negative direct and total effects on the prevalence of melanism across the landscape (Figs. 4, 5; Table 3). This suggests melanic individuals may be avoiding areas with greater road cover, and thus previously observed underrepresentation of the melanic morph among roadkill (Gibbs et al. 2019, Parlin et al. 2025) may be due in part to habitat selection or behavioral avoidance of roads by melanics, rather than being a sole function of differences in visibility to people while driving. Isolating the direct and total effects of road cover while considering confounds (e.g., building cover) provided this new insight into why melanics are underrepresented among roadkill and contributes more evidence that within cities the prevalence of squirrel melanism is constrained by high disturbance.

Another key finding from our models was the strong negative direct effect of predator activity on the prevalence of melanism (Figs. 4, 5; Table 3). A previous translocation experiment showed survival is lower for melanic morph than the gray morph in rural woodlands (Cosentino et al. 2023), and we know the melanic morph is generally more conspicuous than the gray morph (Proctor et al. 2025). As such, crypsis and predation pressure could contribute to patterns in melanism across the urban-rural gradient. This combination of forces might help explain the direct negative effect of predator activity *and* positive total effect of forest fragmentation on melanism. Although we did not find strong evidence of a direct effect of fragmentation on melanism, our analyses support the idea that fragmentation mediates the prevalence of melanism through its negative effect on predator activity (Fig. 5, Supplemental Table S1), leading to a positive total effect of fragmentation on melanism (Fig. 4, Table 3). The melanic morph may benefit from a suppressive effect of fragmentation on predator activity locally, but it is important to note there was a strong positive direct effect of human population density on predator activity (Fig. 5, Supplemental Table S1), indicating predator activity was greatest in the city where melanics are most common. It is possible the melanic morph uses behavioral strategies to mitigate their predation risk in densely populated areas of the city where predator activity is greatest (Sarno et al. 2015; Gaynor et al. 2019; Engel et al. 2020). Such strategies could include different activity patterns in space (including vertically from ground to canopy) and time (diel patterns), resulting in differing susceptibility to predation. Because vertical niche shifts have been linked to competition and predation, climate change, and urbanization (Rankin et al. 2018; Basham and Scheffers 2020; Borden et al. 2021; Gámez and Harris 2022), future research should investigate differences in spatial-temporal behavioral patterns between color morphs and how it may impact the landscape of fear. Additional analyses are also needed to examine the possibility that predator impacts on squirrel melanism vary among predator species.

Despite these novel insights regarding mechanisms contributing to the urban-rural cline in melanism, it is intriguing that melanism is most prevalent in Syracuse yet constrained by components of physical infrastructure that are widespread in cities. This seemingly contradictory pattern may be due in part to the interaction of historic and contemporary forces, including ecological, evolutionary, and social (Des Roches et al. 2021). For example, the melanic morph, once common across the forested landscape, was extirpated in many areas during the period of European colonialism when extensive hunting accompanied deforestation and logging in old growth forests near settlements (Robertson 1973; Cronon 1991; Benson 2013; Thompson et al. 2013). Evidence suggests urban areas may have functioned as refugia for eastern gray squirrels in general, including the melanic morph (Benson 2013; Gibbs et al. 2019). Indeed, changing attitudes about nature led to efforts to deliberately introduce squirrels to cities starting in the late 1800s, celebrating them as “our most loved” species in cities, and establishing a social attitude of tolerance and even admiration (Benson 2013). Despite well-known cases of introductions of the melanic morphs to cities (e.g., Washington DC, Fischman et al. 2021), many squirrel translocations are undocumented. We strongly suspect human-mediated transport and bans on hunting in densely populated cities, including Syracuse, have played important roles in contributing to the maintenance of urban-rural clines in squirrel melanism.

Although such social forces can be difficult to measure, our DAG illustrates it can be important to include unmeasured processes in a causal modeling framework. As just one example, consider the direct negative effect of predator activity on melanism inferred from our analyses. Based on the DAG in Fig. 3B, the adjustment set we used to estimate the direct effect of predator activity on melanism included human population density and fragmentation.

Adjusting for human population density was needed in part to close the backdoor path *Predator activity* ← *Human population density* → *U* → *Squirrel melanism*. If the *Human population density* → *U* → *Squirrel melanism* pathway was *not* included in the DAG, the direct effect of predator activity on squirrel melanism could be estimated without adjusting for human population density. Doing so would lead to a biased estimate of the direct effect of predator activity due to confounding with the unmeasured variable, U. Indeed, when we estimate the direct effect of predator activity on melanism without adjusting for human population density, the estimated effect is no longer negative (posterior mean = 0.04, 95% CI = -0.22, 0.35). This positive bias is due to a positive effect of people on predator activity and the residual positive association between human population density and melanism operating through the unobserved processes. Graphical causal models, such as DAGs, highlight one way in which unobserved processes in urban systems – known or unknown – can be explicitly considered when estimating effects of interest.

A striking demonstration of the value of graphical causal models is seen when comparing estimated effects when appropriately adjusted based on a DAG versus univariate models commonly seen in urban ecology and evolution. For example, had we relied solely on univariate models to estimate the association between squirrel melanism and common urbanization metrics, we might have erroneously concluded that melanism is unrelated to *any* of the measured variables in our DAG other than human population density, as credible intervals for all univariate predictors other than human density overlapped zero (Fig. 4; Table 3). This finding raises significant concerns about using single-variable indices of urbanization. For example, impervious cover is commonly used to represent urbanization (Szulkin et al. 2020), yet our univariate model with impervious cover resulted in a posterior distribution that was largely negative, implying melanism is lowest in areas of high impervious cover (Fig. 4; Table 3). Had we used impervious cover alone as an index of urbanization, we might have concluded there was no evidence for an urban-rural cline, or even weak evidence for a reversed cline where melanism is greater in rural areas. Indeed, our own previous studies using impervious surface as a surrogate for urbanization likely underestimated true clines in melanism (Gibbs et al. 2019; Cosentino and Gibbs 2022). If using single variables as proxies of urbanization to describe biodiversity patterns along urbanization gradients cannot be avoided, our results suggest it is best to use variables that are causal ancestors of environmental change in cities, namely the density of people (Fig. 5).

Distance to city center or composite indices can also be used to describe spatial patterns of biodiversity along urbanization gradients (Moll et al. 2019, Alberti and Wang 2022), but we urge caution about using these variables for causal inference as they often aggregate multiple underlying mechanisms.

## Conclusion

Overall, our research demonstrates that urban impacts on biodiversity are complex and involve a network of causal pathways. Environmental variables commonly used to represent urbanization may at best describe coarse patterns with respect to urbanization effects in ecology and evolution, but they are nearly impossible to interpret mechanistically (Moll et al. 2019). We show that graphical causal models, such as DAGs, can be an effective tool to clarify causal assumptions and inform the design of statistical models to make inferences about a variable’s mechanistic effects. Causal modeling frameworks present great opportunity for disentangling the mechanisms at play in urban systems and understanding the causal drivers of biodiversity within and among cities.

## Acknowledgments

Field assistance was provided by Joelee Tooley and Richard Rich. Access to field sites was provided by the New York Department of Environmental Conservation, City of Syracuse, and select private and municipal landowners.

## Statements and Declarations

### Data Availability

Data and code are available in the following repository: https://github.com/bcosentino/urbanCausalInference

