## Supplemental Materials for "Beyond urbanization metrics: Using graphical causal models to investigate mechanisms in urban ecology and evolution"

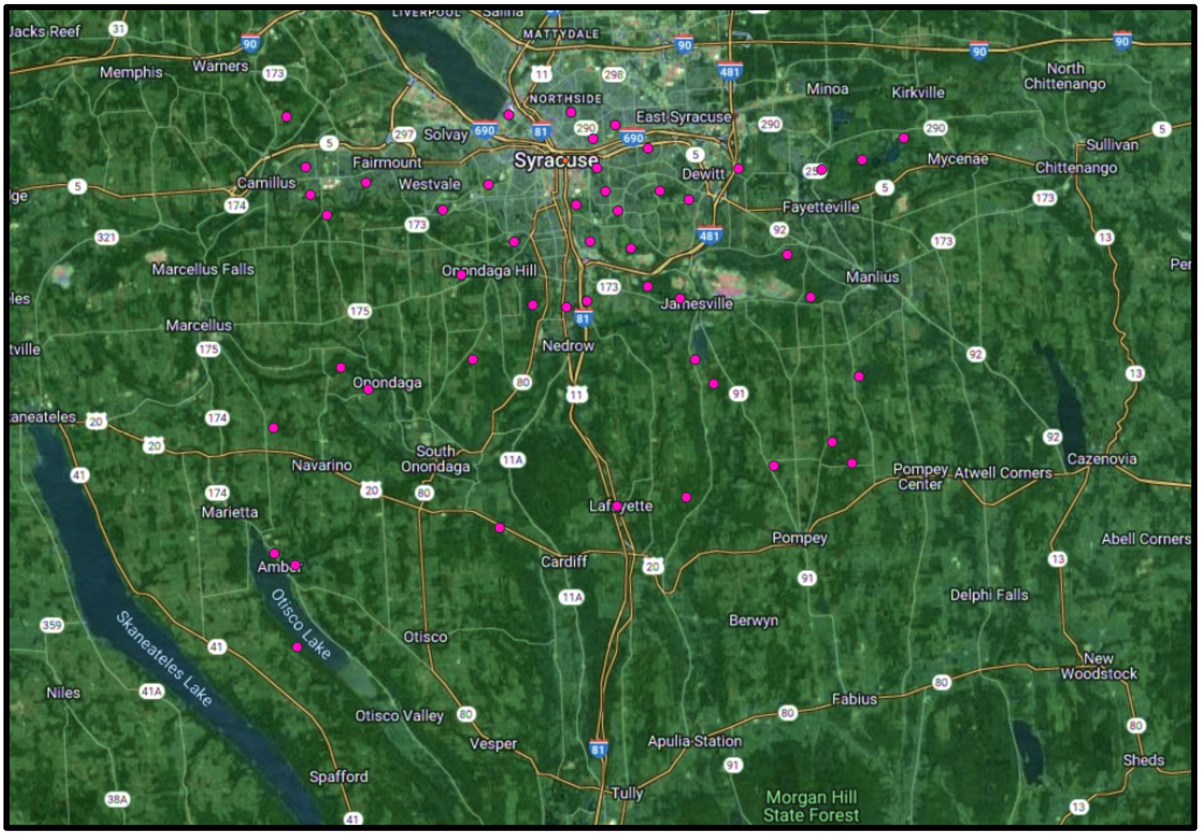


**Supplemental Fig S1.** Map of Syracuse, NY, USA, and the surrounding area. Pink dots represent survey sites across the urban-rural gradient.

**Supplemental Text S1**

We first used generalized linear models with Bayesian inference to examine direct effects among measured variables other than squirrel melanism. For each hypothesized direct effect, we specified a model with an appropriate probability distribution for the response variable, and we included appropriate adjustment covariates to isolate the direct effect of interest based on the assumed causal structures in the DAG (Fig. 3A). We used a beta error distribution for response variables that were proportions (building cover, road cover, impervious cover, forest cover), a lognormal distribution for forest fragmentation, and a binomial distribution for predator activity. Appropriate adjustment variables for each model were determined with the *dagitty* package in R (Textor et al. 2016). Residual plots were visually checked to evaluate model assumptions, and transformations of predictor variables were used when appropriate. We generally used noninformative prior distributions for regression coefficients and variance terms. Predictor variables were standardized to zero mean and unit standard deviation. Models were fit with STAN via the *brms* package in R (Bürkner 2017) with four chains, 2000 warmup iterations, and 1000 final iterations, leaving 4000 total iterations to describe the posterior distribution. We confirmed convergence of parameter estimates by visually examining traceplots and quantifying the Gelman-Rubin statistic (R-hat < 1.01; Gelman and Hill 2007). We summarized the posterior distribution of each regression coefficient of interest with the posterior mean and a 95% credible interval (CI; Supplemental Tab. S1), and we dropped direct effects from the initial DAG (Fig. 3A) when the 95% CI broadly overlapped 0. Based on these initial models, we generated a revised DAG (Fig. 3B) that dropped the following direct effects from the initial DAG: *Human* → *Forest cover*, *Road* → *Predator*, and *Forest cover* → *Predator*.

Next, we examined the hypothesized direct effects on the prevalence of squirrel melanism. We also estimated the total effects of all variables in the DAG, which includes the sum of the direct and indirect effects. Although some variables were not hypothesized to have direct effects on coat color (e.g., impervious cover), they could still have total effects via indirect pathways. In addition to the direct and total effects, we fit univariate models with each measured variable as the sole predictor of melanism to examine how inferences vary between the univariate models and models with appropriate adjustments and estimates of direct and total effects based on the DAG. We used an integrated hierarchical model that combines the camera detection data and visual point count data to estimate the proportion of melanic squirrels at each site (Kéry and Royle 2020; Cosentino et al. 2023). The model included two submodels: 1) A process model to estimate the total squirrel abundance ($N\_total_i$), abundance of each color morph ($N_{ik}$, where $k = 1$ for the melanic morph and $k = 2$ for the gray morph), and proportion melanic ($p\_melanic_i$) at each site $i$; and 2) An observation submodel to estimate individual detection probabilities ($p_{ijk}$) based on repeated surveys $j$ of color morph $k$ at each location $i$. For the observation process, we modeled the counts $y_{ijk}$ of each morph from point count surveys as $y_{ijk}|N_{ik} \sim Binomial(N_{ik}, p_{ijk})$, and we modeled the binary detections $y_{ijk}$ of each morph from trail cameras as $y_{ijk}|N_{ik} \sim Bernoulli(P^*_{ijk})$, where $P^*_{ijk} = 1 - (1 - p_{ijk})^{Nik}$ (Kéry and Royle 2020). Because squirrel activity is influenced by ambient temperature, we modeled individual detection probability as a function of daily ambient temperature (Cosentino et al. 2023):

$$\text{logit}(p_{ijk}) = a0_k + a1_k*\text{temperature}_{ij} + a2_k*\text{temperature}^2_{ij},$$

where for each color morph $k$, $a0_k$ represents the intercept, $a1_k$ represents a linear effect of daily temperature, and $a2_k$ represents a quadratic effect of temperature. A quadratic effect was included as squirrel activity peaks with the intermediate temperatures observed during fall and spring (Cosentino et al. 2023).

For the process submodel, we described the total abundance of squirrels as $N\_total_i \sim Poisson(\lambda_i)$, where $\lambda_i$ is the expected total abundance of squirrels at location $i$. We included a coat color process to describe the abundance of melanic squirrels at each site: $N_{i1} \sim$ Binomial($N\_total_i$,$p\_melanic_i$). We specified linear models of proportion melanic ($p\_melanic_i$) to estimate the direct and total effects of the measured explanatory variables, as well as the univariate models. Each model was fit with a logit link for proportion melanic, the predictor of interest to estimate a direct or total effect, and appropriate adjustment covariates based on the DAG (Fig. 3B). Because total squirrel abundance varies strongly across the urbanization gradient (Cosentino et al. 2023), we included distance to city center as a predictor of abundance ($\lambda_i$) in all models, although exploratory analyses showed that estimates of effects on proportion melanic were not sensitive to the structure of the abundance model.

We used Bayesian estimation to fit the hierarchical models with JAGS 4.3.0 (Plummer 2017) and the package *jagsUI* (Kellner and Meredith 2024) in R (Version 4.2.3, R Core Team 2024). We generally used noninformative prior distributions for intercept and slope parameters (Gelman and Hill 2007). An informative prior was specified for intercepts for individual detection probability and abundance of each color morph because we had prior knowledge of mean detection probability and squirrel abundance (Lemoine 2019; Cosentino et al. 2023). Half-Cauchy priors were used for variance parameters (Gelman 2006). Predictor variables were standardized to zero mean and unit standard deviation. For each model, we ran four chains with enough adaptation, burn-in, and total iterations to achieve convergence of the regression parameters based on visual inspection of the traceplots and confirming R-hat < 1.01. We retained 4000 iterations to describe the posterior distributions.

**Supplemental Table S1.** Posterior means and 95% credible intervals for estimated direct effects among urban covariates based on generalized linear models. Separate models were fit to estimate direct effects of predictor variables on outcome variables with appropriate adjustment covariates based on our directed acyclic graph (Fig. 3A). Appropriate probability distributions (Family) were chosen based on the outcome variable, and transformations were applied as needed to meet model assumptions. Measured predictor variables included human population density (Human), building cover (Building), road cover (Road), impervious cover (Impervious), forest cover (Forest), forest fragmentation (Frag), and mammalian predator activity (Predator).

| Predictor | Outcome | Mean | LCL | UCL | Family | Adjustments |
|---|---|---|---|---|---|---|
| Human (log) | Building | 0.74 | 0.42 | 1.04 | Beta | - |
| Building (logit) | Road | 1.30 | 1.01 | 1.61 | Beta | - |
| Building (logit) | Impervious | 0.77 | 0.31 | 1.26 | Beta | Human, Road |
| Road | Impervious | 0.79 | 0.49 | 1.09 | Beta | Building |
| Human (log) | Impervious | 0.26 | -0.04 | 0.55 | Beta | Building |
| Impervious (logit) | Forest | -0.89 | -1.19 | -0.61 | Beta | Human |
| Human (log) | Forest | 0.03 | -0.21 | 0.29 | Beta | Impervious |
| Impervious (logit) | Frag (log) | 0.28 | 0.03 | 0.52 | Gaussian | Human, Forest |
| Forest | Frag (log) | -0.41 | -0.64 | -0.17 | Gaussian | Human, Impervious |
| Human (log) | Frag (log) | 0.29 | 0.07 | 0.51 | Gaussian | Impervious, Forest |
| Human (log) | Predator | 0.58 | 0.45 | 0.71 | Binomial | Road, Forest, Frag |
| Road | Predator | 0.06 | -0.07 | 0.18 | Binomial | Human, Forest, Frag |
| Forest | Predator | -0.05 | -0.17 | 0.06 | Binomial | Human, Road, Frag |
| Frag (log) | Predator | -0.13 | -0.27 | 0.01 | Binomial | Human, Road, Forest |
